# A linear SARS-CoV-2 DNA vaccine candidate reduces virus shedding in ferrets

**DOI:** 10.1101/2022.09.29.510112

**Authors:** Mathias Martins, Gabriela M. do Nascimento, Antonella Conforti, Jessica C.G. Noll, Joseph A. Impellizeri, Elisa Sanchez, Bettina Wagner, Lucia Lione, Erika Salvatori, Eleonora Pinto, Mirco Compagnone, Brian Viscount, James Hayward, Clay Shorrock, Luigi Aurisicchio, Diego G. Diel

## Abstract

Severe acute respiratory syndrome coronavirus 2 (SARS-CoV-2), the causative agent of coronavirus disease 2019 (COVID-19), has caused more than 600 million cases and over 6 million deaths worldwide. Vaccination has been the main strategy used to contain the spread of the virus, and to avoid hospitalizations and deaths. Currently, there are two mRNA-based and one adenovirus vectored vaccines approved and available for use in the U.S. population. The versatility, low cost and rapid-to-manufacture attributes of DNA vaccines are important advantages over other platforms. However, DNA vaccination must meet higher efficiency levels for use in humans. Importantly, *in vivo* DNA delivery combined with electroporation (EP) has been successfully used in the veterinary field. Here we evaluated the safety, immunogenicity and protective efficacy of a novel linear SARS-CoV-2 DNA vaccine candidate for delivered by intramuscular injection followed by electroporation (Vet-ePorator^™^) in ferrets. The results demonstrated that the linear SARS-CoV-2 DNA vaccine candidate did not cause unexpected side effects, and was able to elicit neutralizing antibodies and T cell responses using a low dose of the linear DNA construct in prime-boost regimen, and significantly reduced shedding of infectious SARS-CoV-2 through oral and nasal secretions in a ferret model.

## 1. Introduction

Severe acute respiratory syndrome coronavirus 2 (SARS-CoV-2) is the viral agent of the deadliest pandemic of the last 100 years. By September 2022, the World Health Organization (WHO) officially registered more than 600 million confirmed cases and over 6 million deaths caused by coronavirus disease 2019 (COVID-19) pandemic worldwide (https://covid19.who.int). The earliest reports of COVID-19 date from December 2019, when health facilities in China reported an outbreak of pneumonia of unknown etiology in Wuhan [1]. Most of the initial cases were linked to the Huanan Seafood Wholesale Market, which sold aquatic animals, live poultries, and several wild animal species [2]. Following the initial reports, human-to-human transmission was confirmed as individuals contracted the infection without ever being at the seafood market [3,4]. To date, the virus RaTG13 isolated from the bat species *Rhinolophus affinis* was found to be the closest related animal coronavirus to SARS-CoV-2, sharing over 96% identity at whole-genome level [5]. However, the genetic distance of approximately 4% (∼1,150 mutations) between RaTG13 and SARS-CoV-2 isolate Wuhan-Hu-1 indicate that transmission likely did not occur directly from bats to humans and further suggest that a yet unidentified animal species may have served as an intermediate host prior to spillover of the virus into humans [6,7].

SARS-CoV-2 belongs to the subgenus *Sarbecovirus*, genus *Betacoronavirus* of the family *Coronaviridae*, and is an enveloped positive-strand RNA virus [8]. The virus encodes a major surface glycoprotein, the spike (S) which mediate receptor binding and is the main target of host immune responses [9]. The S protein mediates SARS-CoV-2 entry into target cells by initially binding to the angiotensin-converting enzyme 2 (ACE2) receptor on the host cell surface (via the S1 domain) and subsequently fusing (via the S2 domain) with host membranes [9–11]. The virus-host receptor interaction occurs between the main functional motif of S, the receptor binding domain (RBD) known as receptor binding motif (RBM), present at the tip of the trimer of viral S protein and the host cell receptor [12,13]. Given its critical role on virus entry, the S RBD is also a major target of host responses to SARS-CoV-2, thus representing a good target for subunit vaccine development.

As of July 15^th^ 2022, there are two mRNA (Pfizer/BNT162b2 and Moderna/mRNA-1273) and one recombinant adenovirus vectored vaccine (Janssen vaccine/Ad26.COV2.S) approved by the U.S. Food and Drug Administration (FDA) for human use in the U.S. [14–16]. Additionally, another 40 vaccine platforms are approved for human use by at least one country. The list includes other non-replicating viral vector vaccines, such as Oxford/AstraZeneca/AZD1222, protein subunit vaccines, including Serum Institute of India/Novavax/COVOVAX and Novavax/Nuvaxovid, one virus-like particle (VLP) vaccine by Medicago/Covifenz, virus inactivated vaccines, such as Sinovac/CoronaVac and Sinopharm (Beijing)/Covilo, and one DNA vaccine, the Zydus Cadila/ZyCoV-D in India [17,18]. The vaccine contains a circular plasmid DNA, which enters the nucleus of the host cells to be transcribed into messenger RNA (mRNA).

Overall, DNA vaccination offers many advantages over other vaccine platforms, such as lack of anti-vector immunity, allowing its use in prime-boost vaccine regimens. In addition, DNA vaccine is rapid-to-manufacture, versatile, relatively easy, inexpensive, and safe compared to the other vaccine platforms. Moreover, the DNA stability at room temperature facilitates the maintenance of this type of vaccine especially in countries or regions with restricted cold-chain resources [19]. Similar to live viruses, DNA vaccines are able to engage both MHC-I and MHC-II pathways, inducing CD8+ and CD4+ T cells [20,21]. However, DNA vaccinations have yet to meet better levels efficiency for use in humans. Several approaches have been explored to improve efficacy of DNA vaccine technologies, including use of improved *in vivo* delivery methods using electroporation (EP) to optimize the cellular uptake of exogeneous DNA [19,22,23]. The EP method applies brief electric pulses to induce reversible cell permeabilization through transient pore formation, from which macromolecules such as DNA can translocate into the intracellular space and deliver the nucleic acid encoding the gene(s) of interest [19].

In the present study we assessed the safety, tolerance, immunogenicity and protective efficacy of a novel linear SARS-CoV-2 DNA vaccine candidate delivered by intramuscular injection followed by electroporation (Vet-ePorator™) using a ferret model of SARS-CoV-2 infection.

## 2. Results

### 2.1 Vaccine safety and tolerance

Twenty-five ferrets (*Mustella putorius furo*) were allocated into 5 groups as follows: sham-immunized (sterile water) (G1), 0.25 mg prime + booster (G2), 1 mg prime + booster (G3), 0.25 mg single dose (G4), and 1 mg single dose (G5). The animals were intramuscularly immunized at day 0 and groups G1, G2 and G3 received a booster dose at day 28 (Figure 1A). The vaccination was immediately followed by intramuscular electroporation. Safety was assessed by monitoring local and systemic adverse reactions for 2 days after each immunization. During the period after the administration of DNA vaccine or placebo (sterile water), there was no significant difference in body temperature between the vaccinated and the sham-immunized groups (Figure 1B). A slight increase in temperature up to 40 ºC was observed one day post the first vaccination and on the booster day in all groups, including sham-immunized animals. These results confirm the safety of the linear DNA vaccination delivered intramuscularly concomitantly by co-localized electroporation (Vet-ePorator™) in ferrets.

**Figure 1.**
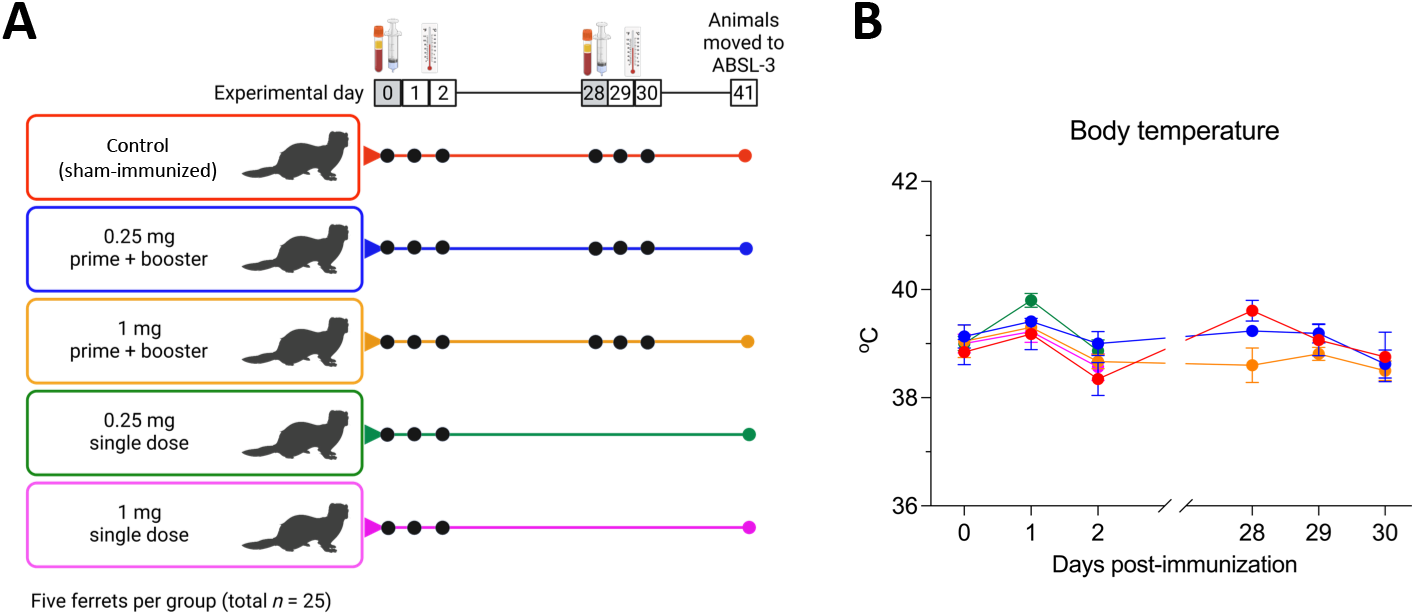
Experimental design, body temperature and body temperature following vaccination with the linear SARS-CoV-2 DNA vaccine candidate. (**A**) A total of twenty-five 12-16-month-old ferrets (*Mustela putorius furo*), with three males and two females per group were allocated into 5 groups, accordingly to the vaccine regimen to be administered: control sham-immunized (G1), 0.25 mg prime + booster (G2), 1 mg prime + booster (G3), 0.25 mg single dose (G4), and 1 mg single dose (G5). (**B**) Body temperature following intramuscular vaccination with the linear SARS-CoV-2 DNA vaccine candidate was measured on the days of vaccine administration, and during the following 2 days as indicted in the graphic.

### 2.2 Vaccine immunogenicity

The serological response to vaccination were assessed using a fluorescent bead-based multiplex and a virus neutralization (VN) assay. Serum samples collected on day 42 post-immunization (pi) were used to assess S-receptor binding domain (RBD)-specific or neutralizing antibodies (NA) levels. All animals from Group 2 (G2) (0.25 mg prime and booster) seroconverted to SARS-CoV-2, as evidenced by detection of antibodies against S-RBD in all 5 immunized animals, and by detection of neutralizing antibodies (NA) in 4 out of 5 animals by day 42 pi (Figure 2). Anti-RBD antibodies were higher in the G2 compared to control group 1 (G1 *p* < 0.01), group 3 (1 mg prime and booster) (G3 *p* < 0.01), and group 4 (0.25 mg single dose) (G4; *p* < 0.05) (Figure 2A). In addition to binding antibodies, 4 out of 5 animals from G2 also presented higher NA on day 42 pi (*p* < 0.05) (Figure 2B), with NA titers of 8 (3 of 4 animals) or 16 (1 of 4). Prior to immunization, all animals were screened and tested negative for SARS-CoV-2 antibodies.

**Figure 2.**
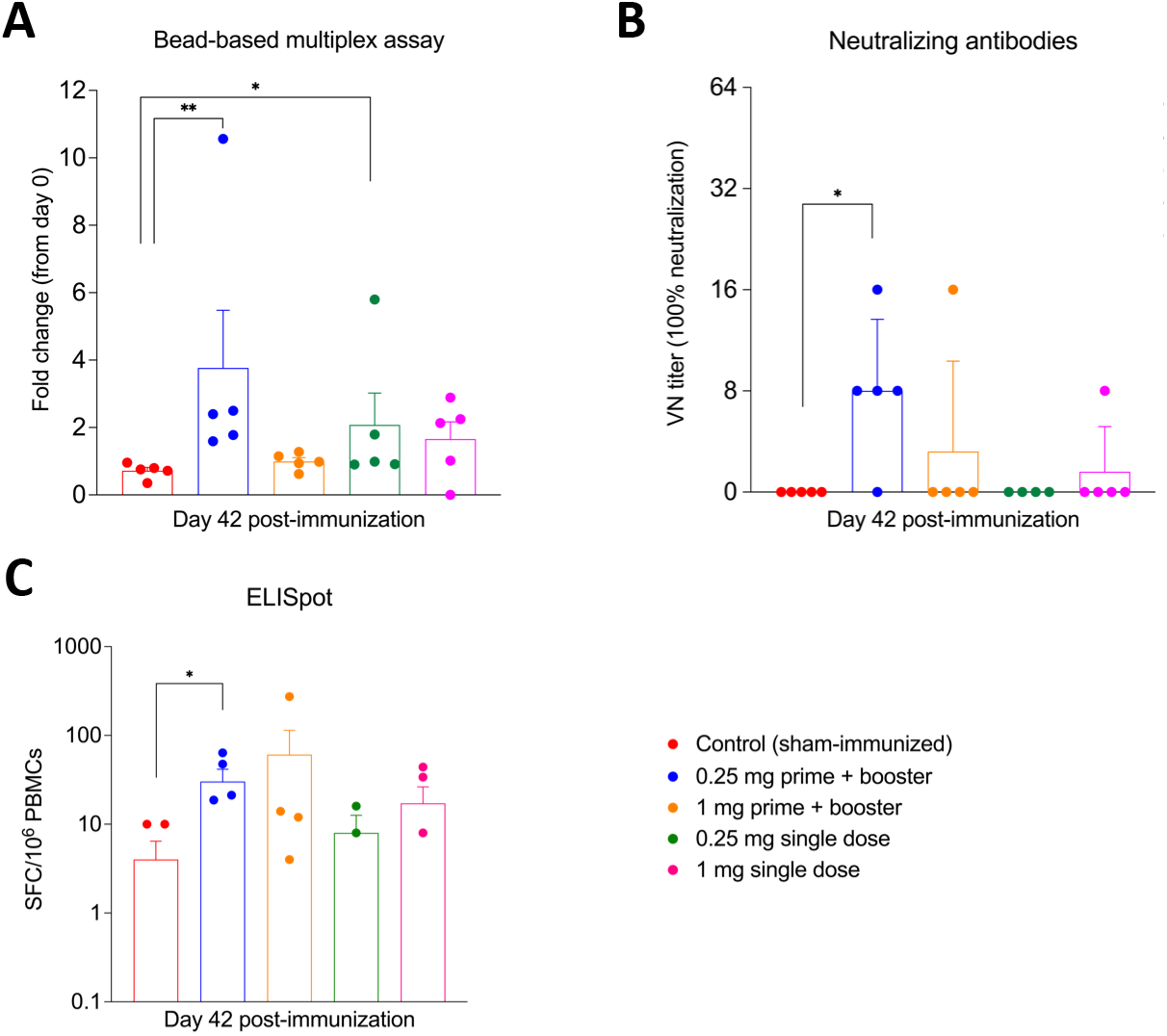
Serological and cellular responses to the linear SARS-CoV-2 DNA vaccine candidate was assessed by bead-based multiplex and virus neutralization assays. (**A**) Antibody responses following immunization were measured bead-based multiplex assay to assess IgG anti-SARS-CoV-2 Spike receptor binding domain (RBD) in serum samples collected on day 42 post-immunization (pi). Results are presented as fold change from day 0 (pre-immunization). (**B**) Neutralizing antibody (NA) responses to SARS-CoV-2 in serum samples collected at day 42 pi. Neutralizing antibody titers represent the reciprocal of the highest dilution of serum that completely inhibited SARS-CoV-2 infection/replication. NA titers are expressed as the reciprocal of serum dilutions presenting 100% neutralization of 100-200 TCID50 of actual SARS-CoV-2 virus. (**C**) SARS-CoV-2-Spike-receptor binding domain (RBD) specific T cell response elicited by linear DNA vaccine was assessed by ELISpot assay. Peripheral blood mononuclear cells (PBMCs) proliferation was measured after stimulated using RBD pool peptides in samples from all ferrets on day 42 post-immunization (pi). * = *p* < 0.05; ** = *p* < 0.01.

T cell responses were examined on day 42, after the booster and before challenge with SARS-CoV-2, and were significantly higher in the G2 vaccinated group in comparison to the sham-immunized group (G1) (*p* < 0.05) (Figure 2C). These results suggest that 0.25 mg of the linear DNA under a prime-boost vaccination schedule was able to induce both humoral and cellular immune responses in ferrets.

### 2.3 Vaccine protection using a heterologous challenge

The vaccine efficacy was assessed using a heterologous virus challenge. On day 42 pi, all control and vaccinated animals were challenged with a SARS-CoV-2 Alpha variant of concern (VOC, isolate NYC853-21). Following the challenge, body temperature, body weight were measured and clinical observations were performed on a daily basis (Figure 3A). No significant differences in body temperature and weight were observed between vaccinated and the control groups. However, a slight increase in body temperature up to 40 ºC on day one and on day six post challenge was noticed in all groups (Figure 3B and C).

**Figure 3.**
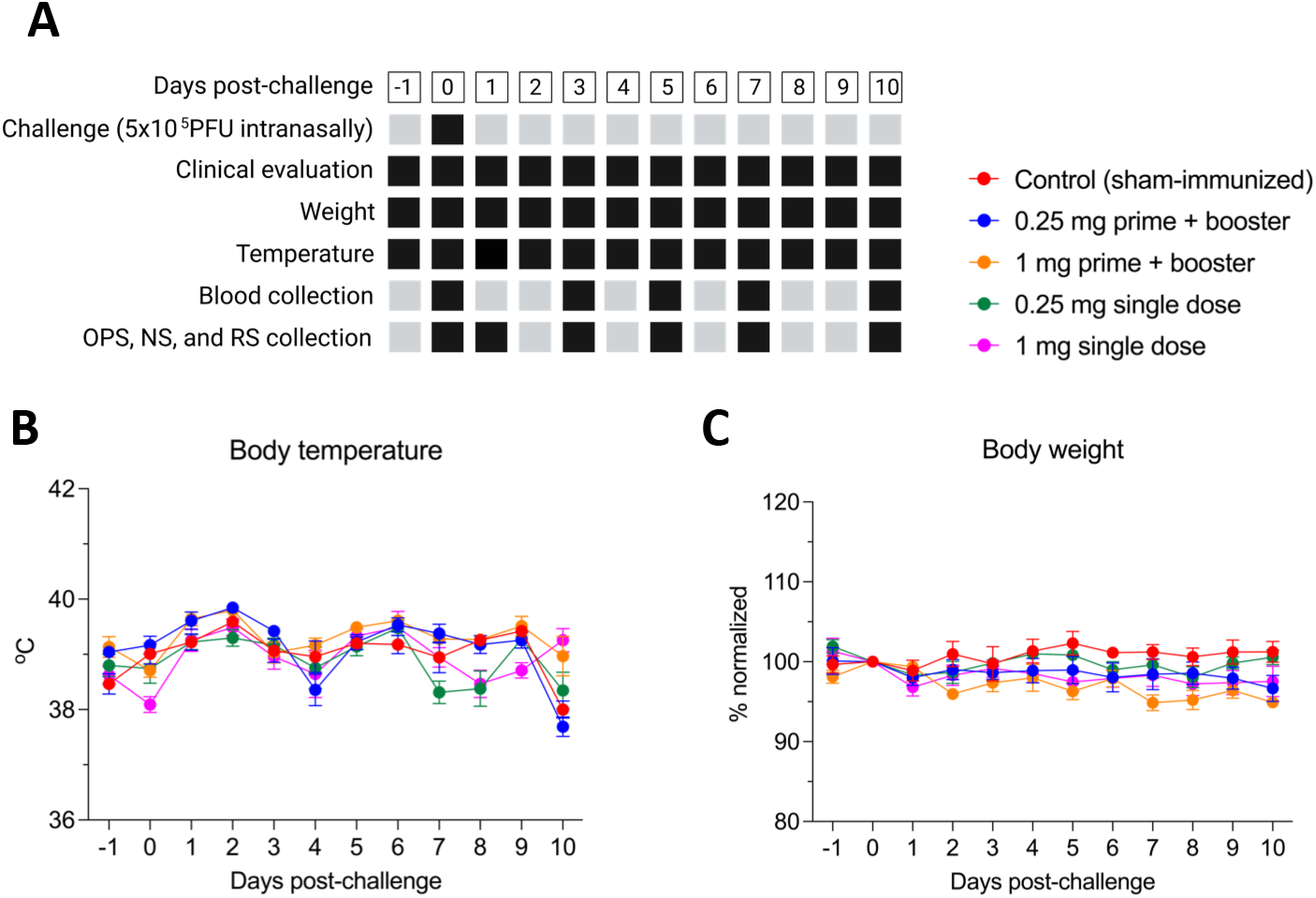
Experimental design, and clinical observations following the intranasal challenge with 5 × 10^5^ PFU of SARS-CoV-2 Alpha variant of concern (isolate NYC853-21). (**A**) Black hatched squares represent actual collection/measure time points for each sample type/parameter described. Clinical parameters, including temperature, body weight, activity, and signs of respiratory disease were monitored daily post-challenge (pc). Oropharyngeal (OPS), nasal (NS), and rectal swab (RS) and blood samples were collected at various times points (black hatched squares). Animals were humanely euthanized on day 10 pc. (**B**) Body temperature and (**C**) body weight following intranasal viral challenge was recorded throughout the experimental period. Data are presented as means + standard error. Body weight was normalized to day 0, which represents 100%.

Following virus challenge, the dynamics of viral replication and RNA shedding was assessed in all animals. Nasal and oropharyngeal secretions and feces obtained through nasal (NS), oropharyngeal (OPS) and rectal swabs (RS) were tested for the presence of SARS-CoV-2 RNA by real-time reverse transcriptase PCR (rRT-PCR). Viral RNA was detected throughout the post challenge (pc) period, between days 1 and 10 pc in all secretions and all groups, regardless of the vaccination regimen used (Figure 4A, B, and C). The highest viral RNA loads were detected between days 1 and 3 pc in oropharyngeal secretions of all groups, which decreased thereafter through day 10 pc. A significant decrease in viral load in oropharyngeal secretions was found only on day 7 for the groups that received 0.25 mg and 1 mg of the DNA vaccine candidate in a prime-boost regimen (G2 and G3) when compared to sham-immunized (G1) (*p* < 0.01) (Figure 4A). Viral RNA load in nasal secretion was reduced on day 7 for G3 in comparison to G5, and on day 10 for G2 (0.25 mg prime + booster) in relation to G5 (*p* < 0.01) (Figure 4B). Overall, shedding of viral RNA in feces was markedly lower than in oropharyngeal and nasal secretions.

**Figure 4.**
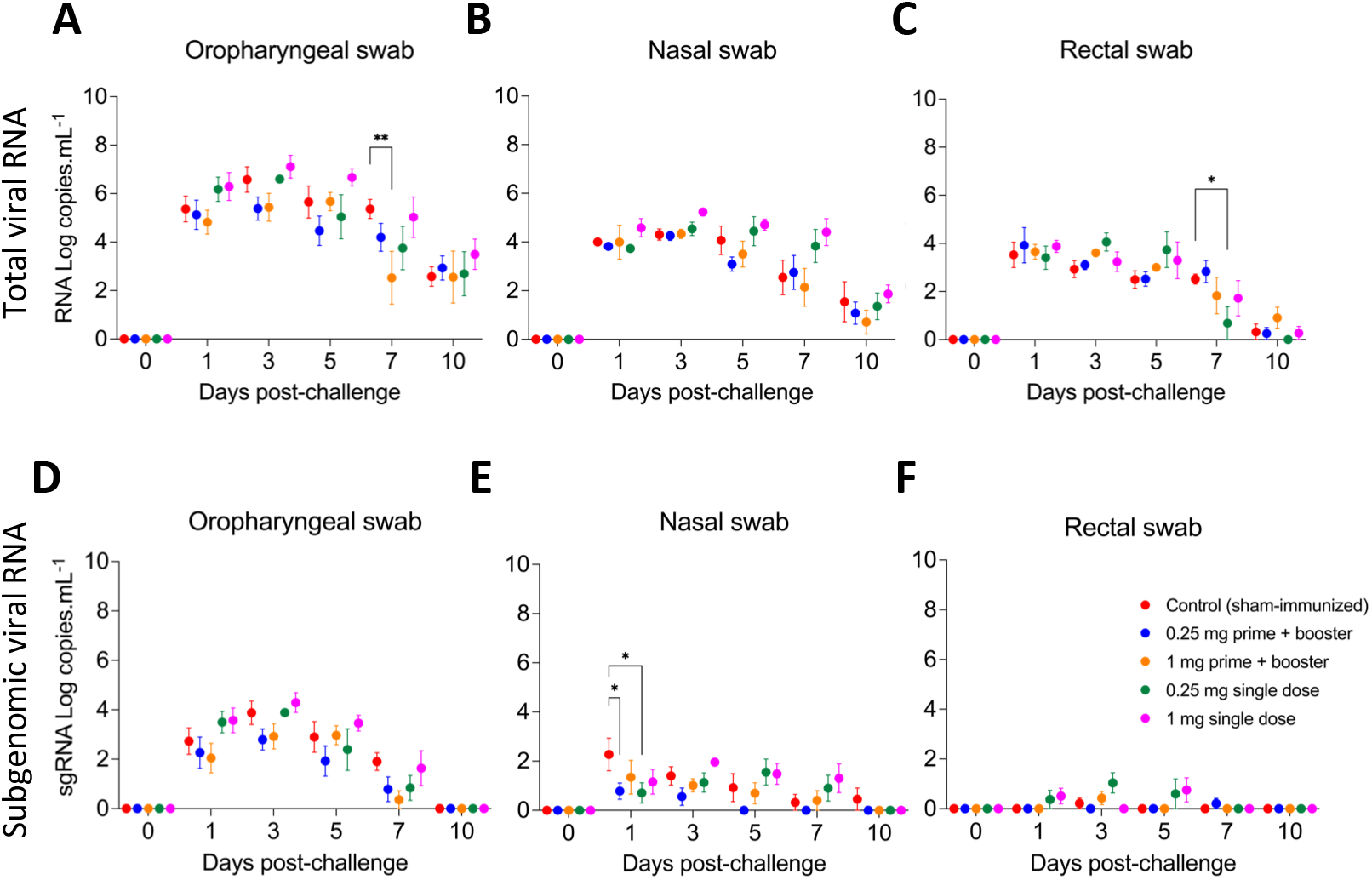
Viral genomic and subgenomic RNA in nasal and oropharyngeal secretions and feces following SARS-CoV-2 challenge. (**A**) Detection of viral RNA in oropharyngeal-(OPS), (**B)** nasal-(NS), and (**C**) rectal swab (RS) samples from ferrets challenged with SARS-CoV-2 Alpha variant of concern. Samples were tested for the presence of SARS-CoV-2 RNA by real-time reverse transcriptase PCR (rRT-PCR) (genomic and subgenomic viral RNA load). (**D**) Detection of subgenomic SARS-CoV-2 RNA (sgRNA) in OPS, (**E**) NS, and (**F**) RS samples of challenged ferrets. Day 0 represents swab samples collected prior to challenge (day 42 post-immunization). Data are presented as means + standard error. * = *p* < 0.05; ** = *p* < 0.01.

In addition to the rRT-PCR for total viral RNA which detects total viral RNA, we also performed RT-PCR for subgenomic RNA (sgRNA). sgRNA was detected from day 1 to day 10 pc in oropharyngeal swabs, peaking on day 3 pc and slowly decreasing until day 7 pc (Figure 4D). A significant decrease of sgRNA in nasal swabs on day 1 pc was detected in both groups that received the 0.25 mg dose (G2 and G4, *p* < 0.05) in relation to sham-immunized ferrets (G1) (Figure 4E). By day 10 pc, only the sham-immunized group still had detectable levels of sgRNA in nasal secretions (Figure 4E). In rectal swabs, the sgRNA was found in very low amounts in all groups (Figure 4F). Although a significant decreased was only found in G2 (0.25 mg prime + booster) and G4 (0.25 mg single dose) related to the sham-immunized group (G1), these results suggest that the ferrets from G2 presented the lowest viral shedding of sgRNA through the bodily secretions tested in our study

In addition to detection of viral RNA, all samples from oropharyngeal and nasal swabs that were positive for SARS-CoV-2 by rRT-PCR were tested for presence of infectious virus. Group 3, which received 1 mg in a prime-boost vaccination regimen, presented a significant decrease in viral loads in oropharyngeal and nasal secretions on day 1 pc (Figure 5A). In addition, while all (5/5) animals from G1 shed infectious virus on day 1 pc, shedding was observed in only three (3/5) and one (1/5) ferret from groups G2 and G3, respectively (Table 1). Notably, after day 1 pc until the end of the study (day 10 pc), no infectious virus was detected from animals from G2 (Figure 5A). As observed in the viral RNA detection in nasal secretions, infectious virus titers were markedly lower in nasal secretions when compared to oropharyngeal samples, with all the vaccinated groups presenting a significant decrease in viral load on days 1 and 3 pc compared to sham-immunized ferrets (Figure 5B). No infectious virus shedding was detected in feces.

**Figure 5.**
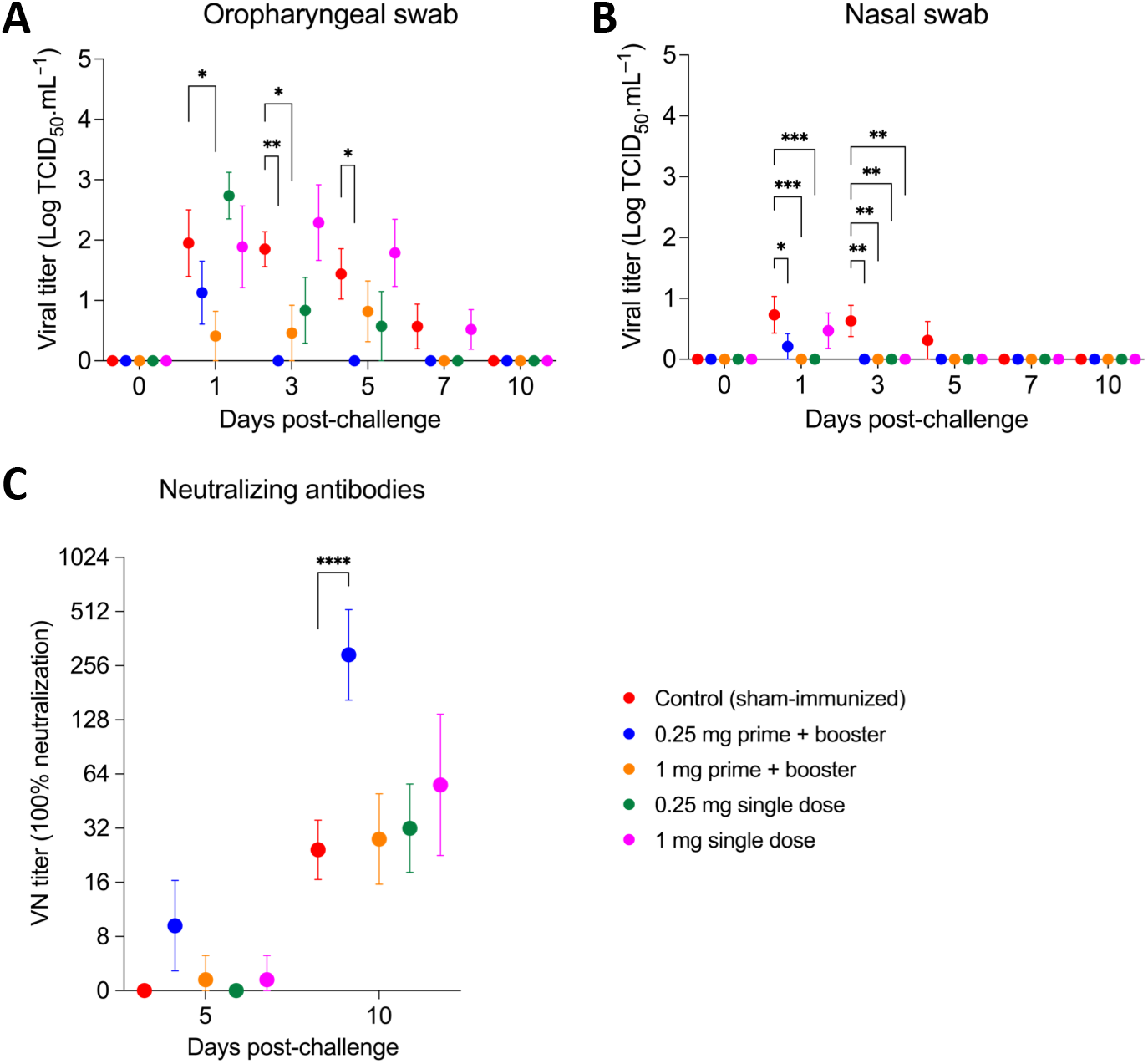
Titer of infectious virus in oropharyngeal secretions and feces and titers of neutralizing antibodies following SARS-CoV-2 challenge. (**A**) Shedding of infectious virus in oropharyngeal, and (**B**) nasal swab samples in ferrets challenged with SARS-CoV-2 Alpha variant of concern. (**C**) Titers of neutralizing antibodies in the animals after SARS-CoV-2 challenge. Day 0 represents swab samples collected prior to challenge (day 42 post-immunization). Data are presented as means + standard error. * = *p* < 0.05; ** = *p* < 0.01; *** = *p* < 0.005; *** = *p* < 0.001.

**Table 1.**
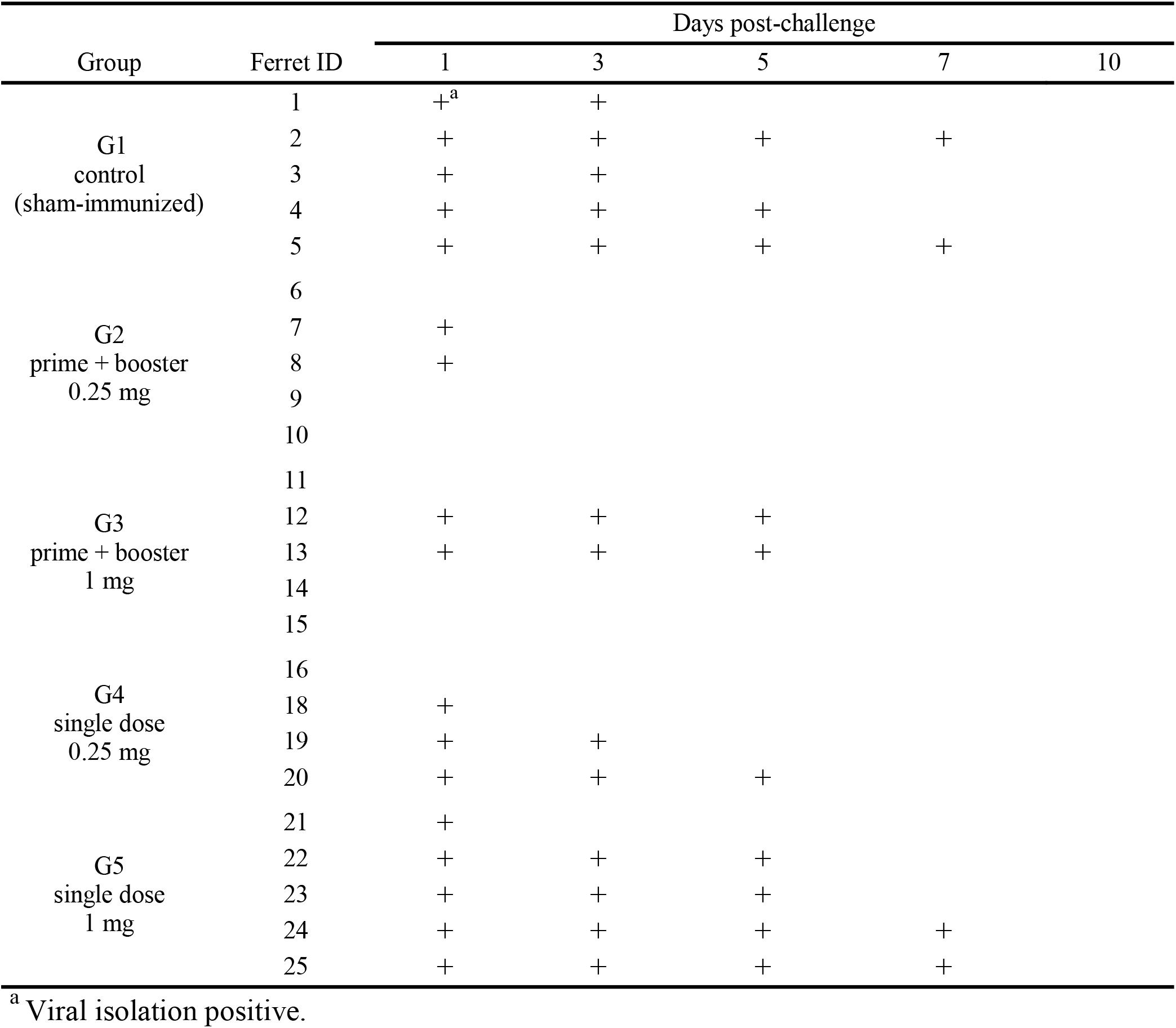
Viral shedding through oropharyngeal secretion of ferrets vaccinated with a linear SARS-CoV-2 DNA vaccine candidate. Animals were intranasally challenge with SARS-CoV-2 on day 0 (day 42 post-immunization) with a SARS-CoV-2 Alpha variant of concern.

All animals seroconverted by day 10 pc (experimental day 52) with NA titers increasing equal or higher than 4x in comparison to the day of challenge (day 0 pc). Interestingly, the highest neutralizing titers were observed in group G2, which was the only group presenting significantly higher NA titers when compared to all other experimental groups (G1, G3-5) (*p* < 0.001) (Figure 5C). These results suggest that 0.25 mg prime-boost vaccination regimen might have been more efficient in priming the immune system, resulting in robust secondary responses to SARS-CoV-2 challenge infection. Together, these results demonstrate that a low dose (0.25 mg) of the linear DNA vaccine administered in a prime-boost vaccination schedule efficiently reduced infectious viral shedding through oropharyngeal and nasal secretions.

## 3. Discussion

DNA-based vaccines are promising platforms due to their potential for rapid and scalable production and manufacturing. Moreover, the stability of DNA at room temperature, makes this platform of particular interest to quickly respond to new emerging pathogens, or to new variants of a circulating agent, as seen with SARS-CoV-2 [24,25]. Of note, DNA uptake by the target cells is a critical step for efficient immunization, which can be significantly enhanced by electroporation [19,26–28].

Ferrets have been used as animal models to study various respiratory viruses including influenza and SARS-CoV-2 [29–32] and to assess pathogenesis, vaccine and drug efficacy and safety, as well as immune responses [31]. In addition to its extensive use to investigate influenza A virus, respiratory syncytial virus and SARS-CoV, the ferret model has been considered a reliable tool to dissect the pathogenesis transmission, and assess therapy and vaccination options for SARS-CoV-2 [29,31,33–35]. Previously, the safety and efficacy of ChAdOx1 nCoV-19 (AstraZeneca) against SARS-CoV-2 challenge was preclinically assessed in ferrets in parallel to non-human primates [36,37]. Here, we used the ferret model to test a linear DNA vaccine candidate against SARS-CoV-2 delivered by electroporation into the epaxial muscle of ferrets.

The two main parts of immune responses against SARS-CoV-2 include cellular and humoral components [38]. Although ideal vaccine aims to achieve sterile infection, those vaccines able to induce neutralizing antibodies and T cell responses will be able to restrain infected cells, restrict viral spread, accelerate viral clearance, and avoid or improve disease outcome [38]. Overall, this linear DNA vaccine not only induced neutralizing antibodies and T cell immune responses, but also reduced viral load in nasal and oral secretions. In addition, the vaccine was shown to be safe with no unexpected adverse reactions following intramuscular vaccination. The minor increase in temperature observed here is a common adverse effect after vaccination against COVID-19 found also in other commercially available vaccines [39,40]. Therefore, there is no evidence of immune-enhanced disease in vaccinated animals found in this study. This has not been the case with all other vaccine studies wherein immunopathologic findings in some vaccination-challenge studies against SARS-CoV have been observed [41–45].

Differences in viral RNA load between control and vaccinated groups were observed only on day 7 post challenge. Most importantly, shedding of infectious virus in oropharyngeal and nasal secretions showed significant reduction from day 1 to 5 pc for the group that received the lowest DNA vaccine dose (0.25 mg) in a prime-boost regimen (G2) in comparison to the unvaccinated animals. Reduced nasal SARS-CoV-2 virus shedding in ferrets was also found in a different preclinical study in which the vaccine tested further became commercially available for use in humans [36]. The difference in protection efficacy between G2 and the other group that received a higher dose of the linear DNA vaccine candidate could be due to a better efficiency of delivery of the linear DNA with the pulse conditions used for electroporation in our study.

In summary, our study shows that intramuscular vaccination with a linear DNA SARS-CoV-2 immediately followed by electroporation (Vet-ePorator™) is safe, elicits both humoral and cellular immune responses with high titers of protective neutralizing antibodies, and significantly reduced shedding of infectious SARS-CoV-2 through oral and nasal secretions in ferrets.

## 4. Material and Methods

### 4.1 Linear DNA Vaccine

Codon optimized complementary DNA (cDNA) encoding the RBD region of SARS-CoV-2 protein S was designed as previously described [24], and chemically synthesized (Genscript, Nanjing, Jiangsu, China). Briefly, synthetic codon-optimized RBD design considering codon usage bias, GC content, CpG dinucleotides content, mRNA secondary structure, cryptic splicing sites, premature PolyA sites, internal chi sites and ribosomal binding sites, negative CpG islands, RNA instability motif (ARE), repeat sequences (direct repeat, reverse repeat, and Dyad repeat) and restriction sites that may interfere with cloning. In addition, to improve translational initiation and performance, Kozak and Shine-Dalgarno Sequences were inserted into the synthetic gene. To increase the efficiency of translational termination, two consecutive stop codons were inserted at the end of cDNA. For the construction of RBD encoding DNA plasmid, the cDNA was amplified via PCR by using sequence-specific primers and directionally cloned into the linearized pTK1A-TPA vector by enzymatic restriction PacI/NotI. The linear DNA amplicon construct encoding RBD was synthetized as previously described [24]. Briefly, for the phosphothioate-modified amplicons, a sulfur atom substitutes the non-bridging oxygen in the phosphate backbone of the oligonucleotide. In addition, the five terminal bases of both the forward and reverse primers were modified to increase DNA amplicon stability. DNA amplification via PCR was performed using a large-scale PCR system using Q5 polymerase from New England BioLabs, USA and resulted in 2780 bp linDNA amplicon expression cassettes. As for purification, the target dsDNA from the PCR amplification was first concentrated by ethanol precipitation and then purified on an Akta Pure 150 FPLC instrument (GE Healthcare, Chicago, IL, USA) with a GE HiPrep 26/60 Sephacryl S-500 High Resolution size exclusion column and 0.3M NaCl running buffer. The purified DNA solution was ethanol precipitated again and resuspended with phosphate buffered saline (PBS) solution to a final concentration of 1 mg.ml^-1^ ± 10% and sterile filtered with a 0.22 μm PES membrane. Further analytical characterization of the linear DNA construct was performed using NanoDrop (Thermo Fisher, Waltham, MA, USA), 2100 Bioanalyzer, and Alliance HPLC System (Waters, Milford, MA USA). Sanger sequencing of the linear DNA construct showed no sequence error as compared to the plasmid DNA template sequence. The linear DNA construct was then lyophilized in a VirTis Genesis Pilot system. All removable internal components of the system were autoclaved. All interior surfaces were wipe sterilized with actril and swabbed for confirmation of sterility by micro testing (plating). The purified sterile linear DNA solution was aseptically filled into the sample vials (2 ml glass amber 15 × 32 mm with 13 mm crimp) at predetermined volume, and a 13 mm 2-leg stoppers were applied to the mouth of each vial. Stoppered vials were then placed into an autoclaved bag and placed in a -20 ± 5 °C freezer for 18 hours preceding lyophilization. After the lyophilization program was run, the vials were stoppered under vacuum, and West 13 mm smooth vial caps were applied and crimped manually. Prior to use in the study, the lyophilized linear DNA constructs were resuspended in 1 ml of sterile water (Hospira Inc., Lake Forest, IL USA).

### 4.2 Ethics statement

The viral isolate used in this study was obtained from residual de-identified diagnostic human nasopharyngeal samples tested at Weill Cornell Medicine kindly provided to us by Dr. Melissa Cushing. The protocols and procedures for transfer of de-identified diagnostic samples were reviewed and approved by the Cornell University Institutional Review Boards (IRB approval numbers 2101010049). The ferrets were handled in accordance with the Animal Welfare Act. The study procedures were reviewed and approved by the Institutional Animal Care and Use Committee at the Cornell University (IACUC approval number 2021-0052).

### 4.3 Animal studies

A total of twenty-five 12-16-month-old ferrets (*Mustela putorius furo*) (three males and two females [*n* = 5] per group were obtained from a commercial breeder (Triple F, Gillett, PA, USA). The ferrets were allocated into 5 groups, as follows: sham-immunized (sterile water) (G1), 0.25 mg prime + booster (G2), 1 mg prime + booster (G3), 0.25 mg single dose (G4), and 1 mg single dose (G5). All animals were housed in the animal biosafety level 2 (ABSL-2) facility at the East Campus Research Facility (ECRF) at Cornell University during the immunization phase. After 72 h acclimation period in the ABSL-2 animals were vaccinated intramuscularly under inhalatory anesthesia using isoflurane with 1 ml injection of the linear DNA vaccine (dose of 0.25 mg for G2 and G4, or 1 mg for G3 and G5) or sterile water (G1, the sham-vaccinated group), followed by intramuscular electroporation (Vet-ePorator™, Carpi, MO, Italy) into the epaxial mid-muscles [28]. The profile of electroporation parameters was checked at every DNA vaccination administration in real time and data were stored into the Vet-ePorator™ archive for each animal. A vaccine booster was administered on day 28 days to animals in groups G1, G2, and G3 following the same procedures described above. For sample collection, ferrets were sedated with dexmedetomidine. Whole blood was collected through cranial vena cava (CVC) puncture using a 3 ml sterile syringe and 23G x 1” needle and transferred into heparin or serum separator tubes on days 0, 28, and 42 post-vaccination. Blood was centrifuged at 1200 x *g* for 10 min and the obtained serum was aliquoted and stored at -20 °C until further analysis. Body weight and temperature were recorded on days 0, 1 and 2 post-immunization (pi) for all the animals, and 28, 29, and 30 pi for animals of groups G1, G2, and G3.

All animals were moved into the ABSL-3 facility at the ECRF at Cornell University on day 39 pi. Following an acclimation period of ∼72 h, on day 42 after prime vaccination, all ferrets were challenged intranasally with 1 ml (0.5 ml per nostril) of a virus suspension containing 5 × 10^5^ PFU of SARS-CoV-2 Alpha variant B.1.1.7 lineage (isolate NYC853-21). All animals were maintained in pairs or individually in Horsfall HEPA-filtered cages, connected to the ABSL-3’s exhaust system. Clinical evaluation was performed daily, including body temperature and body weight measurement, observation of activity level and signs of respiratory disease. Blood, OPS, NS, and RS were collected under sedation (dexmedetomidine) on days 0, 1, 3, 5, 7 and 10 post-challenge (pc), as previously described [29]. All animals were humanely euthanized on day 10 pc.

### 4.4 Serological responses

Antibody responses were assessed using a bead-based multiplex assay. The assay was performed with a few modifications from previously described methods [46]. Briefly, beads were incubated with ferret serum samples diluted 1:200. Antibodies were detected using a biotinylated mouse anti-ferret IgG (H+L). Afterwards, streptavidin-phycoerythrin (Invitrogen, Carlsbad, CA, USA) was added as a final detection step. All incubation steps were performed for 30 min at room temperature and wells were washed after each incubation step. The assay was developed in a Luminex 200 instrument (Luminex Corp., Austin, TX, USA). Assay results for individual animals were expressed as fold change from day 0 (pre-immunization).

Neutralizing antibody (NA) responses to SARS-CoV-2 were assessed by a virus neutralization assay performed under BSL-3 conditions at the Cornell AHDC following protocol established previously [47]. Briefly, twofold serial dilutions (1:8 to 1:1024) of serum samples were incubated with 100 - 200 TCID_50_ of SARS-CoV-2. Following incubation of serum and virus, suspension of Vero E6 cells was added to each well of a 96-well plate and incubated for 48 h at 37 °C, 5% CO_2_ incubator. The cells were fixed and permeabilized as described above and subjected to immunofluorescence assay. Neutralizing antibody titers were expressed as the reciprocal of the highest dilution of serum that completely inhibited SARS-CoV-2 infection/replication.

### 4.5 Cellular responses

RBD-specific T cell response was characterized by ELISpot assay for mouse IFN-_γ_. The assay was performed following manufacturer’s instructions (Mabtech, Nacka Strand, Sweden). An RBD peptide pool composed by 132 out of the 338 peptides covering the whole Spike protein was used for T cell stimulation. Briefly, ELISpot assay was performed by stimulating overnight at 37 °C PBMCs collected on day 42 (after boost and before challenge). The Intracellular Cytokine Staining was performed according to the procedure described previously [24]. PBMCs were stimulated with the RBD peptide pool (5 μg.ml^−1^ final concentration) and brefeldin A (1 μg.ml_−1_ ; BD Pharmingen, San Diego, CA, USA) at 37 °C overnight. DMSO and PMA/IONO (Sigma-Aldrich, Burlington, MA, USA) at 10 g.ml^−1^ were used as internal negative and positive control of the assay, respectively. Spot forming colonies (SFCs) were counted using an automated ELISPOT reader (A.EL.VIS ELIspot reader, Germany). Results are expressed as SFCs/10^6^ PBMCs.

### 4.6 Virus and cells

The Vero-E6/TMPRSS2 (JCRB Cell Bank JCRB1819) were cultured in Dulbecco’s modified eagle medium (DMEM), supplemented with 10% fetal bovine serum (FBS), L-glutamine (2mM), penicillin (100 U.ml^−1^), streptomycin (100 μg.ml^−1^) and gentamycin (50 μg.ml_−1_), and maintained at a 37 °C, 5% CO_2_ incubator. The SARS-CoV-2 Alpha (B.1.1.7 lineage) New York city 853-21 (NYC853-21) isolate used in this study was obtained from residual human anterior nares secretions was propagated in Vero-E6/TMPRSS2 cells. Low passage virus stock (passage 3) was prepared, cleared by centrifugation (2000 x *g* for 15 min) and stored at -80 °C. Genome sequencing was performed to confirm the integrity of the whole genome of the virus stock after amplification in cell culture. Virus titer was determined by plaque assays, calculated according to the Spearman and Karber method, and expressed as plaque-forming units per milliliter (PFU.ml^−1^).

### 4.7 RNA isolation and real-time reverse transcriptase PCR

Ribonucleic acid (RNA) was extracted from 200 μL of cleared swab supernatant of NS, OPS and RS samples. RNA extraction was performed using the MagMax Core extraction kit (Thermo Fisher, Waltham, MA, USA) and the automated KingFisher Flex nucleic acid extractor (Thermo Fisher, Waltham, MA, USA) following the manufacturer’s recommendations.

The real-time reverse transcriptase PCR (rRT-PCR) for total (genomic and subgenomic) viral RNA detection was performed using the EZ-SARS-CoV-2 rRT-PCR assay (Tetracore Inc., Rockville, MD, USA) that targets the virus nucleoprotein (N) gene. An internal inhibition control was included in all reactions. Positive and negative amplification controls were run side-by-side with test samples. For specific subgenomic RNA detection, a rRT-PCR reaction targeting the virus envelope protein (E) gene was used following the primers and protocols previously described [48]. A standard curve using ten-fold serial dilutions from 10^0^ to 10^−8^ of virus suspension containing 10^6^ TCID_50_.ml^-1^ of the SARS-CoV-2 isolate used in the challenge was used for rRT-PCR validation. Based on this standard curve, the relative viral genome copy number in each sample was calculated using GraphPad Prism 9 (GraphPad, La Jolla, CA, USA) and expressed as log10 (genome copy number) per ml.

### 4.8 Virus titration

Samples from OPS, NS, and RS that tested positive for SARS-CoV-2 by rRT-PCR were further subjected to virus isolation and end point titrations under Biosafety Level 3 (BSL-3) conditions at the Animal Health Diagnostic Center (ADHC) Research Suite at Cornell University. For the end point titrations, the samples’ supernatant was subjected to limiting dilutions and inoculated into Vero E6/TMPRSS2 cells cultures prepared 24 h in advance in 96-well plates. At 48 h post-inoculation, cells were fixed and subjected to an immunofluorescence assay (IFA) as previously described [46]. The viral titer at each time point was calculated by using end-point dilutions and the Spearman and Karber’s method and expressed as TCID_50_.ml^−1^.

### 4.9 Statistical analysis and data plotting

Statistical analysis was performed using Mann-Whitney U-tests to compare groups. Statistical analysis and data plotting were performed using the GraphPad Prism software (version 9.0.1). Error bars indicate the standard error of the mean (SEM). Figures 1A and Figure 4A were created with BioRender.com.

## Author Contributions

Conceptualization, Brian Viscount, James Hayward, Luigi Aurisicchio and Diego Diel; Data curation, Gabriela do Nascimento, Antonella Conforti and Bettina Wagner; Formal analysis, Mathias Martins and Antonella Conforti; Funding acquisition, Diego Diel; Investigation, Mathias Martins, Gabriela do Nascimento, Antonella Conforti, Jessica Noll, Elisa Sanchez, Bettina Wagner, Lucia Lione, Erika Salvatori, Eleonora Pinto, Mirco Compagnone, Brian Viscount, James Hayward, Clay Shorrock and Luigi Aurisicchio; Methodology, Mathias Martins, Antonella Conforti, Jessica Noll, Joseph Impellizeri, Elisa Sanchez, Bettina Wagner, Lucia Lione, Erika Salvatori, Eleonora Pinto, Mirco Compagnone, Brian Viscount, Luigi Aurisicchio and Diego Diel; Project administration, Brian Viscount and James Hayward; Resources, Luigi Aurisicchio and Diego Diel; Supervision, Diego Diel; Validation, Mathias Martins and Diego Diel; Visualization, Mathias Martins; Writing – original draft, Mathias Martins and Gabriela do Nascimento; Writing – review & editing, Mathias Martins, Antonella Conforti, Jessica Noll, Joseph Impellizeri, Bettina Wagner, Luigi Aurisicchio and Diego Diel.

## Informed Consent Statement

Not applicable.

## Data Availability Statement

Not applicable.

## Acknowledgments

We thank the Center for Animal Resources and Education (CARE) staff and Cornell Biosafety team for the support and the Animal Health Diagnostic Center at Cornell University for the use of extraction and real-time PCR equipment. We also would like to thank Maureen H. V. Fernandes for her help with euthanasia, and Leonardo C. Caserta for whole-genome sequencing of the challenge virus.

